# AAV-based gene replacement reverses Neurexin-2 downregulation in the cerebellum of a mouse model of phosphomannomutase 2 deficiency (PMM2-CDG)

**DOI:** 10.1101/2025.08.11.669733

**Authors:** M Zhong, K Lai

## Abstract

Phosphomannomutase 2 (PMM2) deficiency is the most common congenital disorders of glycosylation (CDG) with an estimated incidence ranging from 1:20,000 to 1:80,000. Patients manifest a broad spectrum of clinical manifestations, with neurological deficits often emerging as the earliest sign, and may progress to severe multi-organ dysfunction. Mortality reaches 20% by the age of six, primarily due to severe infections, liver insufficiency, or cardiomyopathy. The pathophysiology of the tissue-specific complications remains unclear and there is currently no cure for the disease. In this study, we performed omics analyses of cerebella isolated from a mouse model of PMM2-CDG. RNA-Seq analysis revealed altered gene expression in pathways involved in immune responses and coagulation, while proteomic analysis of proteins enriched by lectin-affinity chromatography identified proteins required for neurodevelopment and neurotransmission. We validated our results by demonstrating significant downregulation of Neurexin-2 in the *Pmm2* knockout (KO) mouse cerebella and showed that its reduced abundance can be reversed by AAV9-*PMM2* gene treatment of the *Pmm2* KO mice.

## Introduction

Congenital Disorders of Glycosylation (CDG) are a group of rare, genetically diverse metabolic disorders resulting from defects in cellular glycosylation pathways^1^. Normal glycosylation is crucial for many cellular functions which include protein folding, cell signaling, hemostasis, and immune responses^2^. Defective glycosylation can thus result in a broad spectrum of clinical manifestations, complicating the diagnosis, management, and treatment of CDG^3^.

Inherited phosphomannomutase 2 (PMM2) deficiency, also known as PMM2-CDG (MIM# 212065), is the most prevalent CDG^4^. PMM2-CDG is inherited in an autosomal-recessive manner and affects approximately 1 in 20,000 to 1 in 80,000 individuals^5-7^. Pathogenic variants in the *PMM2* gene reduce PMM2 enzyme activity, which is essential for the interconversion of mannose-6-phosphate and mannose-1-phosphate, the latter being an important substrate for GDP-mannose synthesis^8^. GDP-mannose deficits can lead to impaired glycosylation of glycoproteins and glycolipids^9^, resulting in PMM2-CDG^10^. By the age of six, about 20% of patients succumb to severe infections, hepatic failure, or cardiomyopathy^5, 11, 12^. Nevertheless, PMM2-CDG is frequently under-diagnosed due to its rather non-specific symptoms^13, 14^. The precise pathophysiology of the disease remains unclear, and there is currently no cure for the disorder.

Earlier, we constructed a mouse model of the disease and demonstrated widespread tissue *Pmm2* knockout (KO) post tamoxifen induction. Moreover, these mice exhibited disease-relevant neurological phenotypes^15^. In this study, we performed transcriptomic and proteomic analyses of cerebella isolated from these animals to delineate the molecular basis of the disease, especially the neurological phenotypes. It should be noted that omics analyses have been performed for other animal/cell models of the PMM2-CDG^16-20^, but our studies here represent the first that combined transcriptomic and proteomic analyses of whole cerebella isolated from a mammalian animal model that has widespread tissue *Pmm2* deletion. In addition, we showed that *in vivo* augmentation of PMM2 activity in our mouse model reversed the downregulation of a protein in the mutant cerebella, further validating our proteomic results.

## Materials and Methods

### Animal model

*Pmm2* KO mice were generated as described previously^15^ and maintained on standard chow post-weaning. Mice were housed in standard laboratory cages within temperature (22°C-23°C)- and humidity (30%-70%)-controlled rooms under a 12:12 light: dark cycle. Genotyping was performed on all mice following a previously published protocol^15^. All procedures conducted with the animals were approved and performed in full compliance with the guidelines outlined in the Guide for the Care and Use of Laboratory Animals and were approved by the University of Utah Institutional Animal Care and Use Committee.

### AAV9-based *PMM2* expression vector

AAV9-*PMM2* was designed by the Lai Lab and synthesized at Vector-Builder Inc (IL, USA) on a fee-for-service basis. A dose of 7.5*10^13^ viral genomes (VG) per kg body weight (BW) was systemically administered *via* the tail vein using our previously published technique^15^.

### Western blot analysis

For Western blotting of total proteins, tissue samples from *Pmm2* KO and control mice were homogenized in RIPA buffer (50 mM Tris HCl, pH 7.2; 150 mM NaCl; 0.1% NP-40; 0.5% sodium deoxycholate; 0.1% SDS; 1 mM EDTA) supplemented with one Pierce Protease and Phosphatase Inhibitor Mini Tablet (1 tablet per 10 mL; Thermo Fisher, #A32963), 1 mM sodium orthovanadate, and 20 mM sodium fluoride, freshly prepared. Samples were subjected to 8% SDS-PAGE and transferred to a nitrocellulose membrane (#1620097, BIO-RAD, CA, USA) using established protocols from our laboratory^21^. For Western blot analysis, Neurexin-2 was initially detected by primary antibody against NRXN2 (#ab34245, Abcam, MA, USA) until it was later discontinued by the vendor at which point, we began to use anti-pan-Neurexin antibody (#EPR27983-60, Abcam, MA USA) and the identification of Neurexin-2 from other neurexins was based on its unique size. Antibodies against α-Tubulin (Proteintech, #11224-1-AP, IL, USA), and β-Actin (#CST-3700S, Cell Signaling Technology, MA, USA) were supplied by Proteintech and Cell Signaling Technology, respectively. Detection of primary antibodies was performed using secondary antibodies (#926-32211, #926-32350, LI-COR, NE, USA). We adhered to the recommended titers for all antibodies by the respective vendors. All immunoblots were visualized using the Odyssey DLx Imager (LI-COR, NE, USA). Quantitative analysis of fluorescence signals was conducted using Image J software, with results normalized to the corresponding β-Actin or α-Tubulin levels in the same samples.

### RNA isolation, library preparation, and sequencing

Cerebella from three control (*Pmm2* ^+/+^ *Cre*^*+/-*^) and *Pmm2* KO (*Pmm2* ^fl/fl^ *Cre*^*+/-*^) mice were collected two months after tamoxifen induction, flash-frozen, and sent to the Biorepository and Molecular Pathology Shared Resource at Huntsman Cancer Institute for total RNA isolation. RNA sequencing was then conducted at the University of Utah Huntsman Cancer Institute High-Throughput Genomics Core Facility. RNA concentration was assessed using a Qubit RNA BR Assay Kit (Fisher Scientific, Cat. No. Q10211), and RNA quality was evaluated with an Agilent Technologies RNA ScreenTape Assay (Agilent Technologies, CA). Total RNA samples (100-500 ng) were hybridized with Ribo-Zero Gold to effectively deplete cytoplasmic and mitochondrial rRNA. Stranded RNA sequencing libraries were prepared using the Illumina TruSeq Stranded Total RNA Library Prep Gold kit (Illumina, Inc., CA) with TruSeq RNA UD Indexes (Illumina, Inc., CA). Purified libraries were assessed for quality on an Agilent Technologies 2200 TapeStation with a D1000 ScreenTape assay (Agilent Technologies, CA). The molarity of adapter-modified molecules was determined by quantitative PCR using the Kapa Biosystems Kapa Library Quant Kit (Agilent Technologies, CA). Individual libraries were normalized to 1.30 nM in preparation for Illumina sequencing analysis. Sequencing libraries were chemically denatured and loaded onto an Illumina NovaSeq flow cell using the NovaSeq XP workflow (Illumina, Inc., CA). Once transferred to an Illumina NovaSeq 6000 instrument, a 150-cycle paired-end sequencing run was conducted using the NovaSeq 6000 S4 reagent Kit v1.5 (Illumina, Inc., CA).

### Analysis of RNA-Seq data

For differential expression analysis, mouse GRCm38 FASTA and GTF files were downloaded from Ensembl release 112, and a reference database was created using STAR version 2.7.9a, with splice junctions optimized for 100 bp reads^22^. Optical duplicates were removed from the paired-end FASTQ files using BBMap’s Clumpify utility (v38.34), and adapter trimming was performed using Cutadapt 1.16. The trimmed reads were aligned to the reference database with STAR, and mapped reads were assigned to annotated genes in the GTF file using featureCounts version 1.6.3. Output files from Cutadapt, FastQC, STAR, and featureCounts were summarized using MultiQC to identify sample outliers. Read counts were then normalized with the DESeq2 analysis package, and a regularized log transformation was applied to manage variance among low-abundance transcripts, enabling comparison of transcript variation across all expression levels. Differentially expressed genes were identified with a q-value threshold of > 0.05. Differentially expressed genes in the contrasts were analyzed using GSEA version 4.2.2 to identify enriched gene sets in the RNA-Seq data compared to curated gene sets from the hallmark and canonical pathways of the Molecular Signatures Database. A false discovery rate threshold of < 0.05 was applied to determine significantly enriched pathways.

### Proteomics sample preparation

Six cerebella from control (*Pmm2* ^+/+^ *Cre*^*+/-*^) and *Pmm2* KO (*Pmm2* ^fl/fl^ *Cre*^*+/-*^) mice were harvested two months after tamoxifen induction. Glycoproteins were isolated using the Pierce™ Glycoprotein Isolation Kit with WGA (Thermo Fisher Scientific™, Waltham, MA) and sent to the proteomics core facility at the University of Utah. The protein was acetone-precipitated, and total protein concentration was quantified using the Pierce™ BCA Protein Assay Kit. For de-glycosylation, 10 μg of proteins were treated with the denaturing Protein Deglycosylation Mix II (New England Biolabs, Ipswich, MA) following the manufacturer’s protocol. The proteins were reduced in 1x S-TRAP buffer (50 mM TEAB, pH 8.5, 5% SDS) with 20 mM DTT at 37 °C for 30 minutes, alkylated with 40 mM IAA at room temperature for 45 minutes in the dark, and acidified with 55% phosphoric acid to achieve a final concentration of 4.5%. Protein digestion was performed using Protifi S-TRAP micro columns (Protifi, Fairport, NY), with 10 μg of protein digested by 0.75 μg of trypsin/LysC for 3 hours at 47°C. The peptides were eluted from the column, dried completely, and resuspended in 300 μL of 0.1% TFA. All samples were desalted using Pierce™ Peptide Desalting Spin Columns as per the manufacturer’s instructions and then resuspended at 0.1 μg/μL in 0.1% formic acid for LC-MS/MS analysis.

### LC-MS/MS Mass Spectrometry and data analysis

Reversed-phase nano-LC-MS/MS was conducted on a nanoElute 2 (Bruker, Billerica, MA) coupled with a Bruker timsTOF Pro2 mass spectrometer featuring a nanoelectrospray source. A total of 200 ng from each sample was injected onto a ReproSil C18 150 mm x 0.15 mm nanocolumn (Bruker, Billerica, MA) maintained at 50°C. Peptides were eluted using a gradient of reversed-phase buffers: Buffer A (0.1% formic acid in 100% water) and Buffer B (0.1% formic acid in 100% acetonitrile) at a flow rate of 0.5 μL/min for a 45-minute run. The gradient started at 5% Buffer B, increasing to 28% over 40 minutes, and was held at 95% Buffer B for the final 3 minutes.

The timsTOF Pro2 was operated in PASEF data-dependent acquisition MS/MS scan mode to generate the custom peptide library. The TIMS section was operated with a 120 ms ramp time at a rate of 7.93 Hz and an ion mobility scan range of 0.6-1.4 V·s/cm2. MS and MS/MS spectra were recorded from 100 to 1,700 m/z. A polygon filter was applied to select against singly charged ions. The quadrupole isolation width was set to 3 Da. The mass spectrometer was operated in PASEF data-independent acquisition MS/MS scan mode to analyze each experimental sample. The TIMS section was operated with a 120 ms ramp time at a rate of 7.93 Hz and an ion mobility scan range of 0.6-1.4 V·s/cm2. dia-PASEF window parameters were set to mass width of 25 Da and 47 mass steps/ cycle. MS and MS/MS spectra were recorded from 147 to 1,322.6 m/z. The custom spectral library was created using FragPipe v20.0 software against the uniprot_ref_mouse database (downloaded on 12-7-2023 with 17,191 proteins).

Protein abundances based on peak intensity were calculated using DIA-NN software version 1.8.1, allowing for one missed cleavage after trypsin/LysC digestion. No fixed modifications were considered, while variable modifications included methionine oxidation and cysteine carbamidomethylation (IAA), with a mass tolerance of 15 ppm for precursor ions and 10 ppm for fragment ions. Results were filtered with a false discovery rate of 0.01 for both proteins and peptides.

### Statistical analyses

Statistical analyses were performed using SPSS 22 and GraphPad Prism 10. Continuous variables were presented as means with standard deviations or medians with interquartile ranges (IQRs). The Shapiro-Wilk test evaluated the normality of quantitative data. For normally distributed data, the t-test or analysis of variance (ANOVA) was used as appropriate; for non-normally distributed data, the Mann-Whitney U test or Kruskal-Wallis H test was applied as suitable. A two-sided p-value of < 0.05 was regarded as statistically significant.

## Results

### Transcriptomic analysis of cerebella from a mouse model for PMM2-CDG revealed differential gene expression in multiple vital processes including coagulation and cell adhesion

We examined cohorts of three mutant and three control mice two months post-tamoxifen induction, utilizing RNA sequencing to assess transcriptional changes in *Pmm2* KO mice. We identified 13 differentially expressed genes (DEGs), with five upregulated and eight downregulated (false discovery rate < 0.05). To evaluate the biological significance of these DEGs, we performed functional enrichment analysis using KEGG, GO, and Reactome. Key immune responses, including allograft rejection, antigen processing and presentation, graft-versus-host disease, type I diabetes mellitus, autoimmune thyroid disease, viral myocarditis, and systemic lupus erythematosus, were significantly affected. Dysregulation was also observed in genes involved in cell adhesion and endocytosis, as well as development-related pathways such as Hedgehog signaling and those implicated in basal cell carcinoma. Notable changes were detected in hemostatic and detoxification mechanisms mediated by complement and coagulation cascades, along with cytochrome P450 enzymes, and disruptions in cysteine, methionine, and selenoamino acid metabolism. Additionally, transcriptional changes were evident in circadian rhythm regulation and asthma-related signaling pathways (**Fig. 1**).

**Fig. 1.**
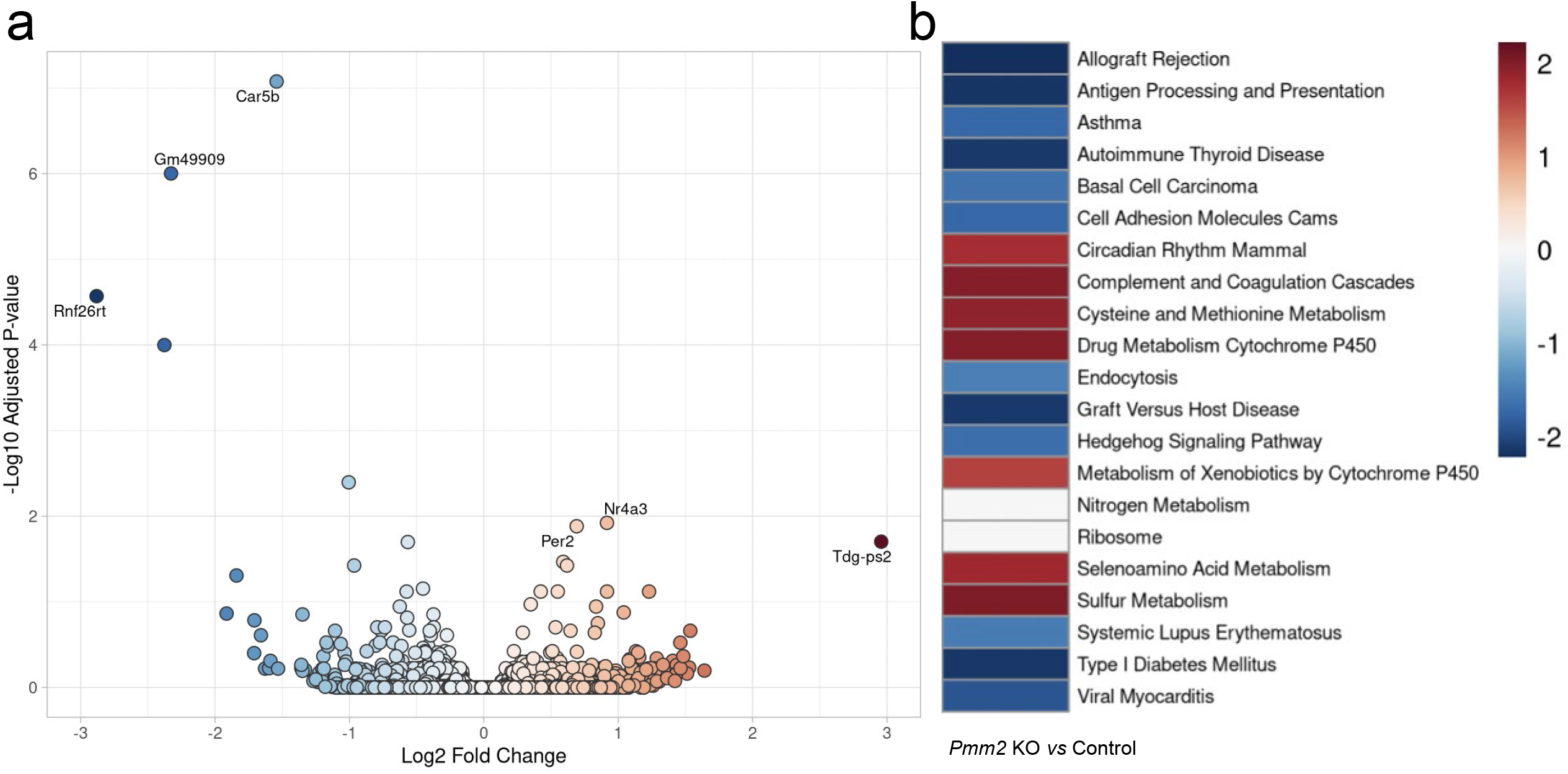
Transcriptomic analysis of cerebella from *Pmm2* KO and control mice. (a) Volcano plot depicting differentially expressed genes in the cerebella of *Pmm2* KO mice (b) KEGG functional enrichment analysis highlighting the biological significance of these differentially expressed genes.

### Proteomic analysis of cerebella from a mouse model for PMM2-CDG identified unique protein changes in neurodevelopment and neurotransmission

We performed proteomic analysis of Wheat germ agglutinin (WGA)-affinity purified proteins isolated from the cerebella of six *Pmm2* KO mice and six control mice, detecting a total of 2,466 proteins. The volcano plot (**Fig. 2a**) illustrated the distribution of proteins, revealing notable changes in global protein levels in *Pmm2* KO cerebella. To validate our results, we showed that Neurexin-2, one of the differentially expressed glycoproteins detected, was downregulated in these samples upon immunoblot analyses (**Fig. 2b**). To evaluate the functional impact of global protein changes, we performed Reactome enrichment analysis, identifying significant disruptions in pathways essential for neuronal development and signaling. Notably, affected processes included neuronal system development, neurotransmitter receptor function, chemical synaptic transmission, second messenger cascades, axon guidance, WNT signaling, G-protein-coupled receptor signaling, insulin receptor signaling, and the metabolism of amine-derived hormones.

**Fig. 2.**
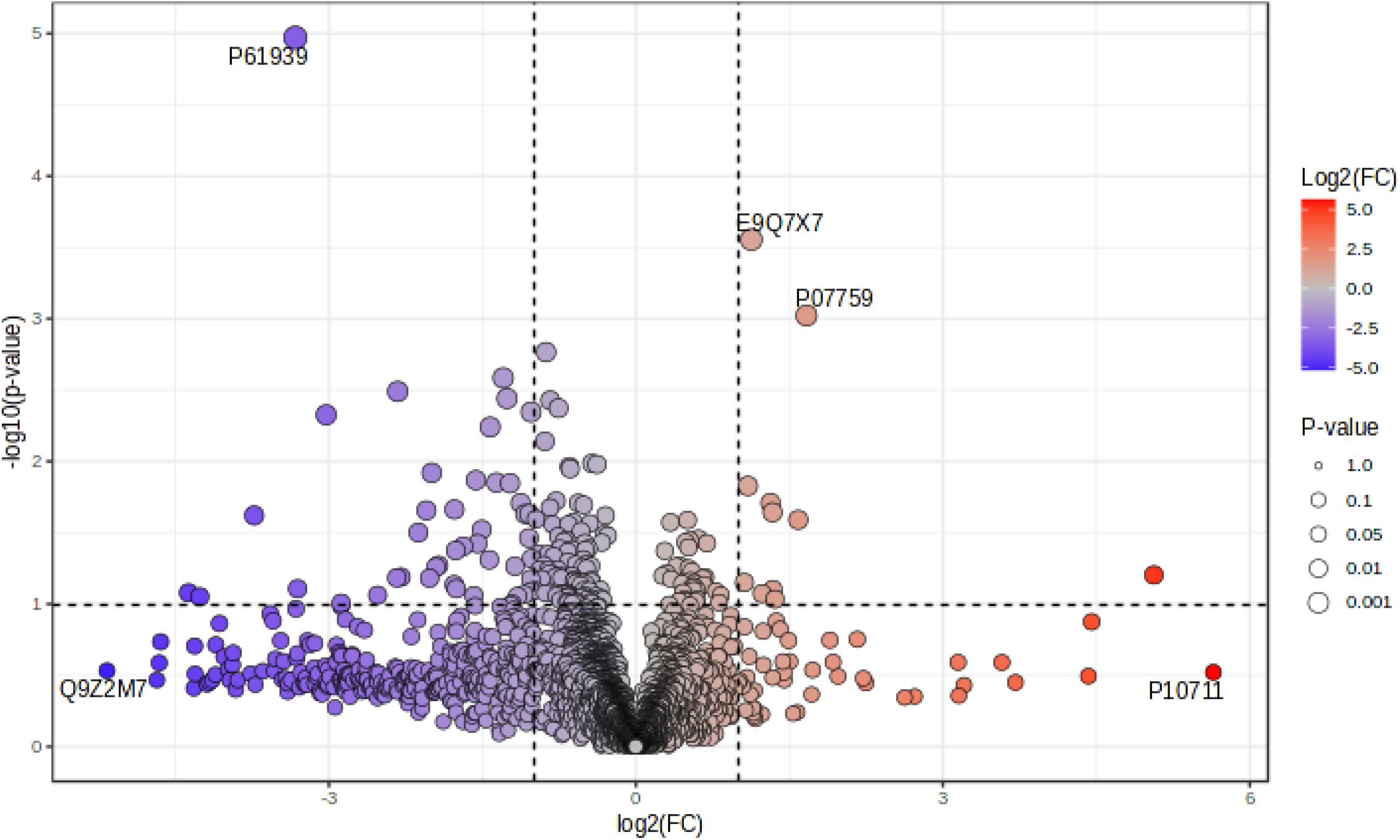
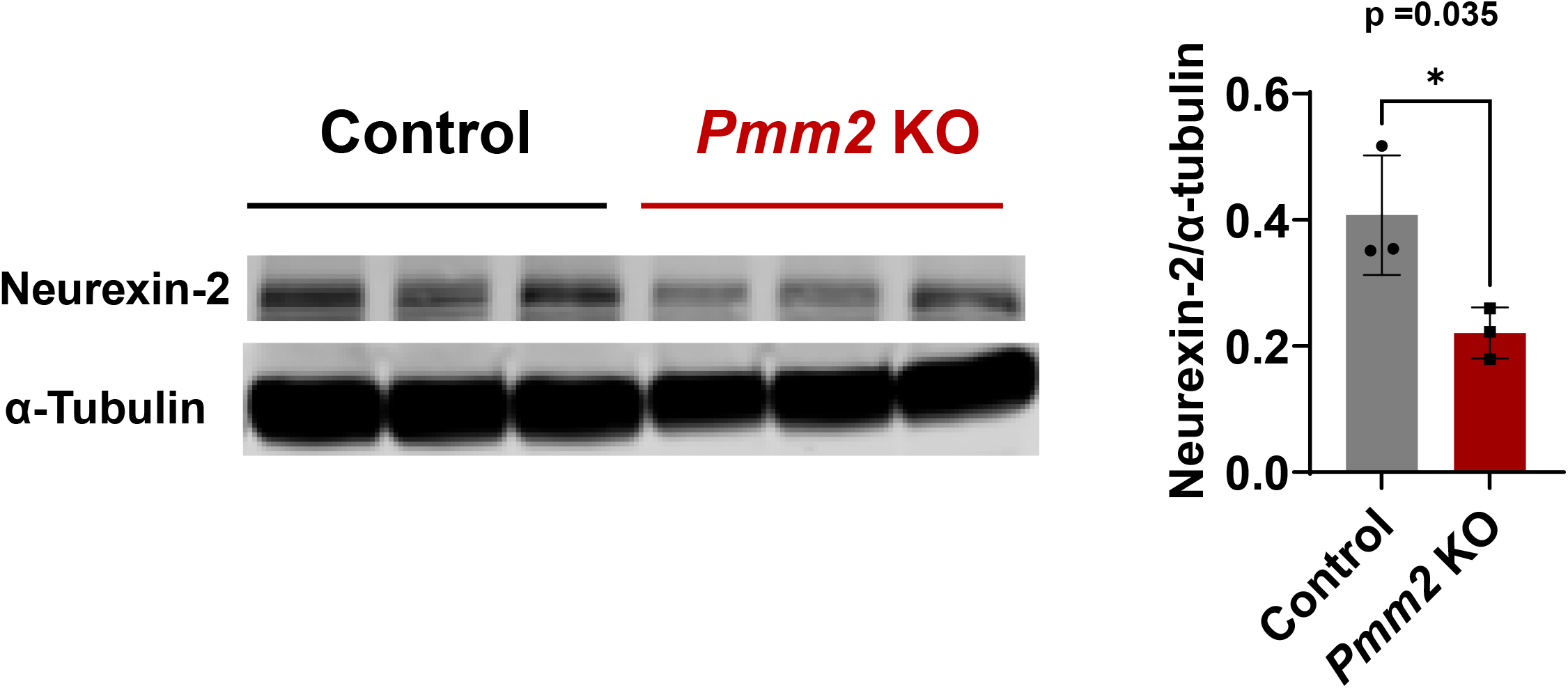
Proteomic analysis of cerebella from *Pmm2* KO and control mice. **(a)** Volcano plot illustrating the differentially expressed proteins in the cerebella of *Pmm2* KO mice. **(b)** Immunoblot analysis of Neurexin-2 protein expression in the cerebella, quantified relative to α-Tubulin as a loading control. *, p < 0.05.

### AAV9-*PMM2* gene replacement therapy reversed the Neurexin-2 down-regulation in the treated *Pmm2* KO mice

To test the hypothesis that augmentation of *PMM2* expression would reverse abnormal (glyco)proteomic changes in PMM2-CDG, we examined Neurexin-2 deficiency in *Pmm2* KO mice treated with AAV9-*PMM2* gene replacement. We administered AAV9-*PMM2* (7.5*10^13^ vg/kg) *via* tail vein injection to the cohort of six *Pmm2*^fl/fl^ *Cre*^+/-^ mice at three weeks of age, prior to tamoxifen induction one week later. Western blot analysis revealed enhanced Neurexin-2 expression in the treated *Pmm2* KO mice compared to the untreated group **(Fig. 3a)**. To test the efficacy of AAV9-*PMM2* gene replacement therapy in a more clinically relevant context, we administered AAV9-*PMM2* (7.5*10^13^ vg/kg) *via* tail vein injection to the cohort of three *Pmm2*^fl/fl^ *Cre*^+/-^ mice at six weeks of age, 10 days post tamoxifen induction, which corresponded to approximately 11.5 years old in human^23^. Western blot analysis showed increased Neurexin-2 expression in the treated *Pmm2* KO mice compared to the untreated group **(Fig. 3b)**.

**Fig. 3.**
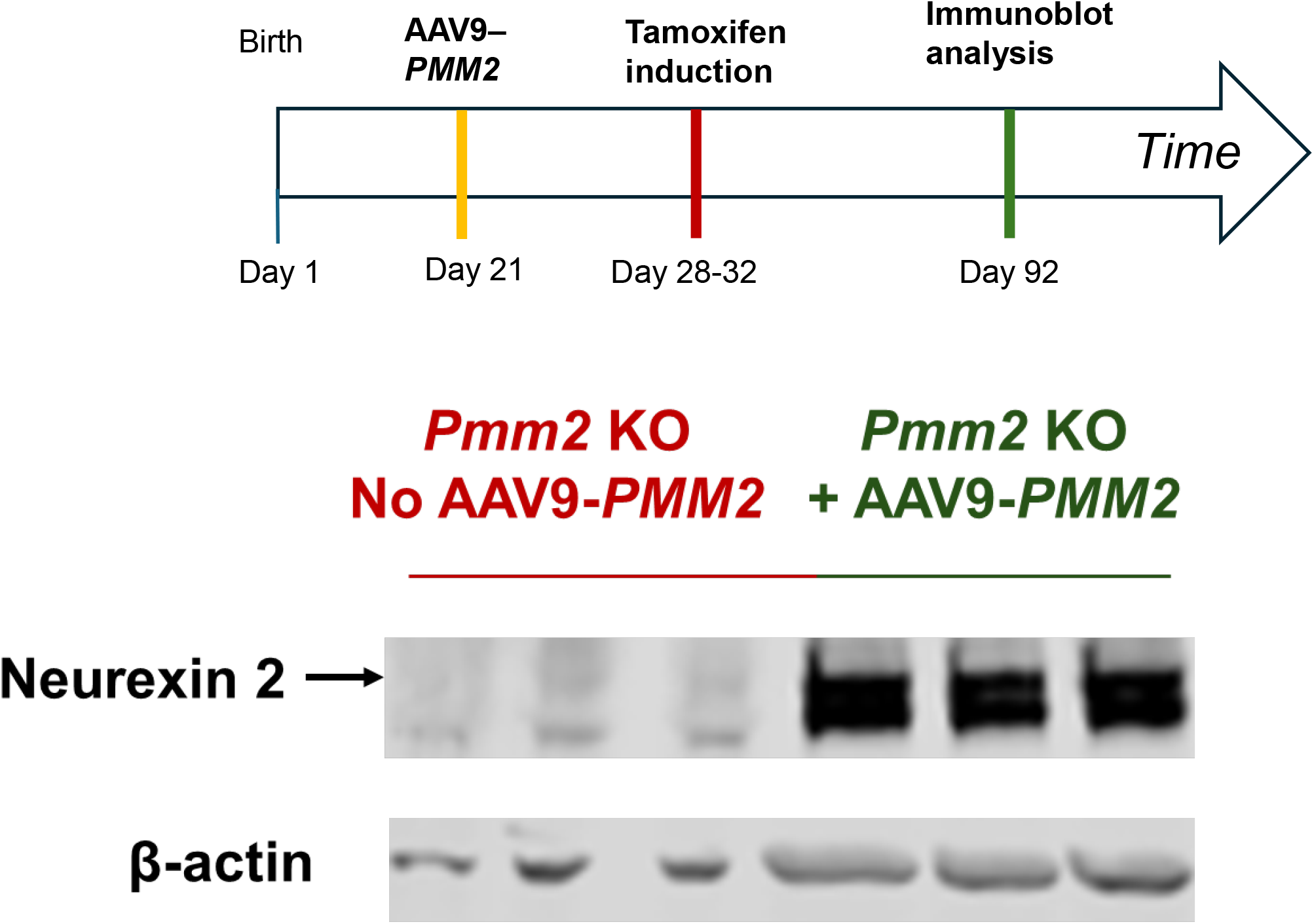
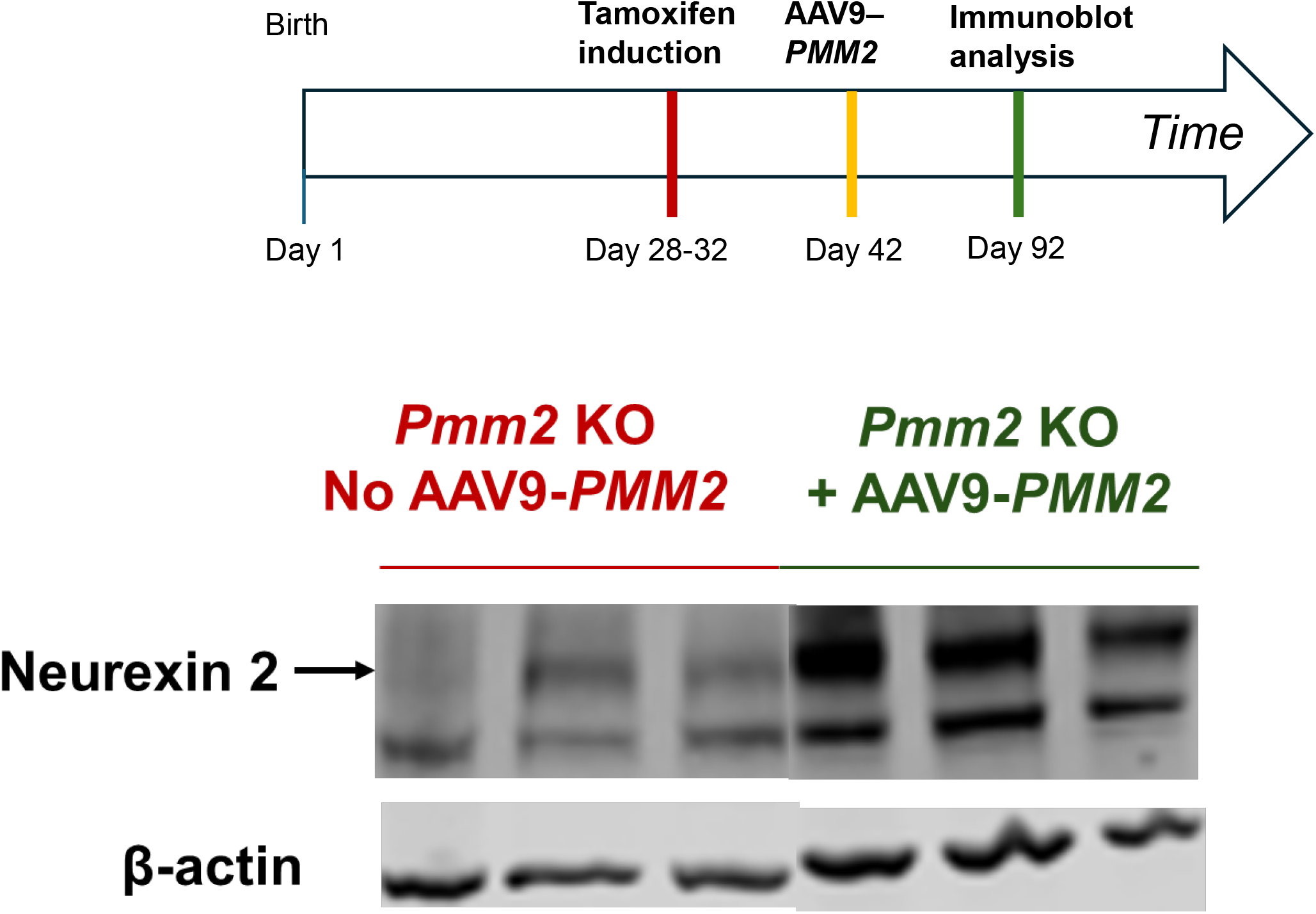
AAV9-*PMM2* gene replacement therapy reversed the Neurexin-2 downregulation in the treated *Pmm2* KO mice. **(a)** Experimental design overview and immunoblot analysis of Neurexin-2 expression in *Pmm2* KO mice with AAV9-*PMM2* treatment one week before tamoxifen induction. **(b)** Overview of the experimental design and immunoblot analysis of Neurexin-2 expression in *Pmm2* KO mice treated with AAV9-*PMM2* 10 days after tamoxifen induction.

## Discussion

With an estimated prevalence of 1:20,000 in certain studied populations^6^, PMM2-CDG is the most common CDG. Patients with PMM2-CDG present with variable features ranging from isolated neurologic involvement to severe multi-organ dysfunction^24^, which makes it difficult for the early diagnosis. Severe cases frequently result in mortality within the initial years of life, with a global mortality rate as high as 20% during this critical period. Yet, the precise pathobiology of different tissue-specific complications remains largely unknown.

In this study, we utilized a novel tamoxifen-inducible mouse model of PMM2-CDG to dissect the molecular cascades of events leading from cellular Pmm2 deficiency to the neurological complications. Although it is not the first mouse model of the disease, many of the models constructed before did not live long enough for the desired studies. Our unique conditional *Pmm2* KO nature of our model allows the animal to survive and grow for the described work. Like many well-characterized conditional gene knockout mouse models, we induced *Pmm2* KO postnatally. As a result, we conceded that our model offered limited mechanistic insights into the role of *Pmm2* in normal embryonic development. However, we believe that it remains a valuable model for PMM2-CDG, mainly because one cannot attribute the pathophysiology of the disease solely to prenatal PMM2 deficit. After all, the metabolic needs for PMM2 does not stop after a baby is born since synthesis of glycoproteins and glycolipids continues throughout postnatal life, and postnatal PMM2 deficiency could be as detrimental, if not more, than the potential prenatal insults. In fact, our preliminary characterization of this novel mouse model at phenotypic and molecular levels confirmed that it recapitulates many patient phenotypes^15^. These results support the notion that the pathophysiology of the disease cannot be *solely* caused by prenatal PMM2 deficiency, or we will not see any disease-related phenotypes in our *Pmm2* KO mice when induction of *Pmm2* KO took place after birth. Last but not least, we prefer to adopt a holistic approach in our studies and therefore, we used a mouse model that has widespread tissue Pmm2 deficiency so that it can mimic what happens in patients best.

When we began our transcriptomic analyses, we focused on the cerebella of the animals as cerebellum is a common target organ in PMM2-CDG. Interestingly, we identified pathways that implicate key immune responses, which include allograft rejection, antigen processing and presentation, graft-versus-host disease, type I diabetes mellitus, autoimmune thyroid disease, and systemic lupus erythematosus that were significantly affected in the Pmm2-deficient cerebella (**Fig. 1**). These findings are both significant and relevant given the immunocompromised state of many surviving patients. As PMM2-CDG is a glycosylation disorder, it is not too surprising that dysregulation was also observed in genes involved in cell adhesion and endocytosis, as well as development-related pathways such as Hedgehog signaling and those implicated in basal cell carcinoma (**Fig. 1**). But what surprised us more was the changes detected in hemostatic and detoxification mechanisms mediated by complement and coagulation cascades (**Fig. 1**). Again, these changes are highly relevant given the coagulopathy experienced by many patients. If we had not employed a holistic approach to use the entire organ to perform these studies, pathways like the coagulation pathway altered in endothelial cells in the cerebella would not have been identified at all. This experience exemplified the importance of performing -omics analyses in the entire organ, as well as single cell populations.

As we proceeded to proteomic analysis, we opted to employ WGA-affinity chromatography to enrich glycoproteins for the analyses. We chose this procedure not only because we want to focus on glycoproteins in our studies but also to validate the technique as a cost-effective alternative for investigators who do not have access to the more advanced instrumentation and expertise required for glycoproteomics. In our studies, we identified many glycoproteins implicated in neurodevelopment and neurotransmission being affected in the Pmm2-deficient cerebella (**Fig. 2a**). We further validated our techniques and approach by performing immunoblot analyses of a protein called Neurexin-2 (**Fig. 2b**). We chose this protein because of its roles in neurotransmission and potential implications in the pathophysiology of the disease. Its significance in synaptic transmission makes it a compelling target for investigation.

Neurexin-2, encoded by the *NRXN2* gene, is a neuronal cell surface protein in the neurexin family. It is a critical cell adhesion molecule and a receptor for synapse formation and maintenance in the central nervous system^25^. Neurexin-2 has multiple isoforms, primarily alpha and beta, generated through alternative promoter usage and splicing^26, 27^. These isoforms are essential for the precise alignment of synaptic components, modulation of calcium channel activity, and maintenance of the balance between excitatory and inhibitory neurotransmission^28^. Increasing evidence indicates that mutations or altered expression of Neurexin-2 can disrupt synaptic function and contribute to various neurological disorders. In fact, it has been proposed as a potential biomarker for the neurodegenerative disorder Alzheimer’s disease (AD)^29, 30^. Its reduced expression correlates with synaptic loss and cognitive decline, hallmarks of AD progression^31^. In this study, we showed significant down-regulation of Neurexin-2 in *Pmm2* KO mice compared to controls. Notably, we reversed the downregulated cerebellar Neurexin-2 levels through AAV9-*PMM2* gene replacement therapy in *Pmm2* KO mice. These results suggested that Neurexin-2 may serve as a novel biomarker for the management of PMM2-CDG treatment by gene replacement therapy.

Despite the novel discoveries made in this study, it has limitations. First, since our proteomic analysis used WGA-affinity enriched glycoproteins, most non-glycosylated proteins encoded in the altered transcriptomic data are unlikely to be identified unless they bind tightly to glycoproteins enriched by WGA affinity chromatography. We plan to identify these proteins when we repeat the proteomic analysis without the lectin chromatography steps in the future. Moreover, while we validated our proteomic results and the WGA affinity enrichment through immunoblot analyses of Neurexin-2, we did not go further to reveal the molecular cascades that link Pmm2 deficiency to reduced abundance of Neurexin-2. Future work will focus more on mechanistic studies that will improve our understanding of the pathophysiology of the disease. Lastly, we only conducted our studies at a single time point, which cannot capture the entire disease development process. Upcoming studies will include more time points so that we can better understand the progression of the disease.

## Data Availability

All data needed to evaluate the conclusions in the manuscript are included.

## Acknowledgements

We want to acknowledge the scientific and technical support, which include the use of equipment and instrument from the following University of Utah Core Facilities:

(1) University of Utah Mutagenesis Generation and Detection Core Facility (Crystal Davey, PhD)

(2) University of Utah Transgenic and Gene Targeting Mouse Core Facility (Crystal Davey, PhD)

(3) University of Utah Huntsman Cancer Institute (HCI) High-Throughput Genomics Core Facility (Brian K. Dalley, PhD)

(4) HCI Bioinformatic Analysis Core Facility (David Nix, PhD; Qing Li, PhD)

(5) Research reported (RNA isolation) in this publication utilized the Biorepository and Molecular Pathology Shared Resource at Huntsman Cancer Institute at the University of Utah and was supported by the National Cancer Institute of the National Institutes of Health under Award Number P30CA042014.

(6) Proteomics analysis was performed at the Mass Spectrometry and Proteomics Core Facility at the University of Utah. Mass spectrometry equipment was obtained through internal shared resources fund.

## Author Contributions

MZ conceived the study, performed experiments, analyzed data, and wrote the manuscript. KL obtained funding, conceived the study, and edited the manuscript. Both authors approve the final version of the manuscript.

## Ethics Approval

Animal care and usage was approved by the University of Utah Institutional Animal Care and Use Committee (Protocol # 2350).

